# Supervised Automation of Cell Counting in Confocal Microscopic Cochlear Imaging Datasets Using Macro in Imaris

**DOI:** 10.1101/2024.07.24.604970

**Authors:** Muhammad Taifur Rahman, Nashwaan Ali Khan, Mobin Ibne Mokbul, Ibrahim Razu, Shakila Mahmuda Fatima, Samia Sultana Lira, Cristina Garcia, Peter Eckard, Marlan R Hansen

## Abstract

Analyzing confocal microscopic data of biological samples using the Imaris software poses challenges due to its time-consuming nature involving tedious multiple steps and possibility of human errors. Here, we developed a supervised automation protocol to minimize manual input in cell and spot counting on confocal images obtained from mouse cochlear sections. The protocol increases efficiency by incorporating image recognition and object-oriented macros. Moreover, the protocol being adaptable allows scientists in diverse other fields to customize it for their specific needs.

**Graphical Abstract:** 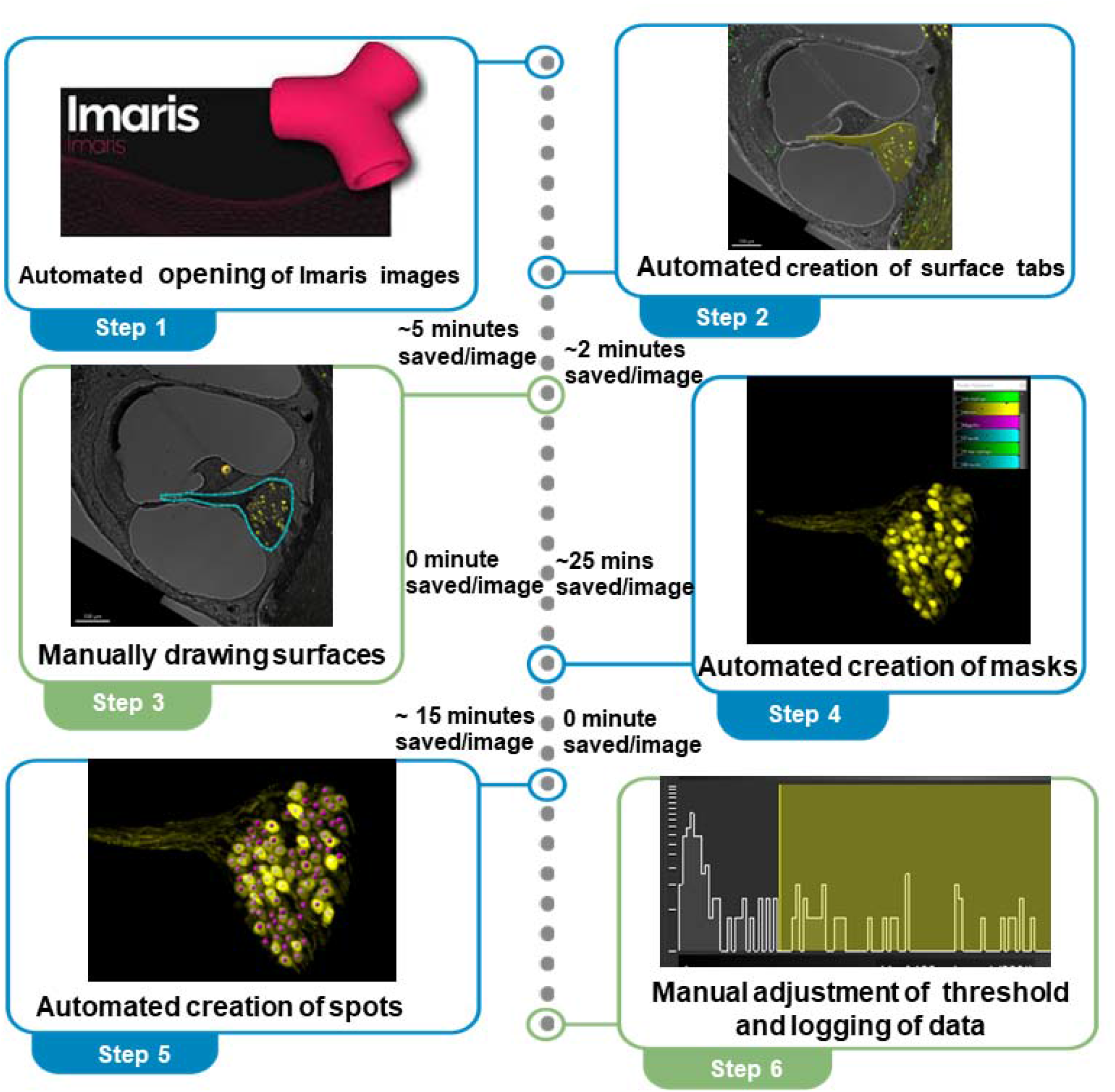

## Introduction

Microscopy is an inevitable technique in research in life science and biomedical sciences with application in medical diagnosis. The last decade has witnessed a flourish in the development of microscopy technologies allowing researchers to analyze anatomy of biological specimens in more detail and facilitating insights into complex processes. Conventional microscopies have technical limitations as it can acquire images only for a thin-cut section of tissue, thereby restricting capability to analyze thick tissue samples as well as *in vivo* investigations (Sanderson, 2020). Developed by Marvin Minsky in 1955, Confocal Microscopy (CM) addresses these limitations in optical imaging providing images with high resolution and of more contrast in comparison to the conventional microscopy (Elliott, 2020). Additionally, it allows researcher to acquire images inside live cells and tissues and to track dynamic events including cell migration and blood flow. Moreover, the imaging technique can perform combination of images for multiple sections resulting in a virtual 3-dimensional image, which is similar to reconstructing computer tomography or magnetic resonance image of the tissue sample (Nwaneshiudu et al., 2012).

Due to its capacity to detect minute structures, CM has become a useful tool for studying cochlear anatomy, such as synapses of the auditory nerve onto the hair cells of the organ of Corti and neurons of the spiral ganglion. On top of that, flexibility in using immunocytochemical staining (e.g., Hoechst labelling), fluorescent reporter (e.g., GFP, YFP) as well as antibodies in detection of numerous cell types (e.g. neurons, macrophage) via CM makes it highly versatile for wide range of applications (Hardie, MacDonald and Rubel, 2004).

For analyzing CM images, commercially available software (e.g., Imaris, Zeiss ZEN, Nikon NIS, HCImage) and open-source software (e.g., ImageJ, Cellprofiler/Cell Analyst, Neuronstudio) are used. Imaris, a software developed by Oxford Instruments is one of the most popular software packages for image analysis due to the wide array of manipulation and analysis it allows. However, it involves time consuming and laborious repetitive processes for image analysis tasks. With a goal to increase efficiency and productivity, we aimed to develop a protocol for utilizing macros in Imaris software for confocal image analysis using mouse cochlear samples. This newly developed supervised semi-automated protocol not only can speed up optical image analysis but also is amenable to using for different biological specimens.

### Preparation for the Manual Processing

Installation of the necessary software: For the manual processing, the requirements are a suitable computer with a Windows operating system (64-bit) and Imaris 10.1 software [Oxford Instruments] available at https://imaris.oxinst.com/learning/view/article/imaris-10.1-trainable-ai-image-analysis-for-everyone.

**Note:** The user can use any image processing software as preferred. Here, we describe Imaris 10.1 Software as a reference.

**Table 1:**
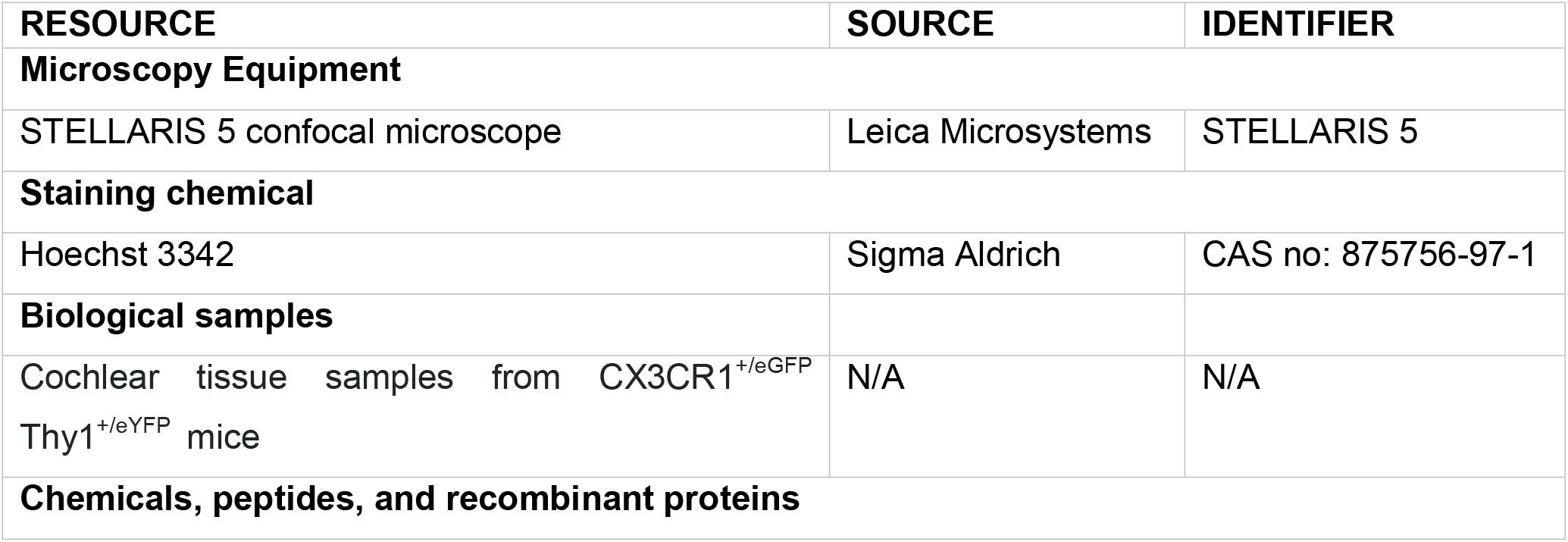
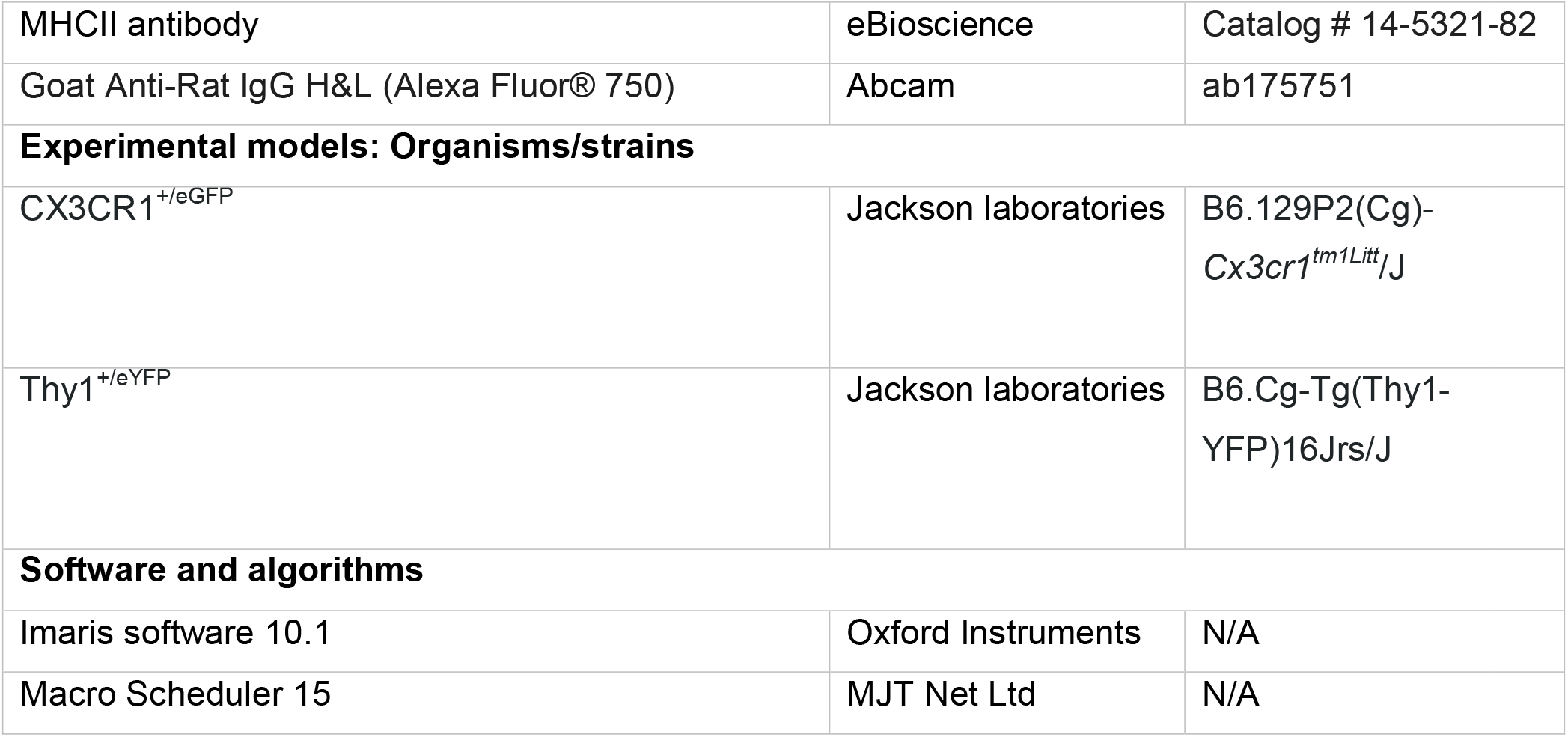
Resources and equipment used in the protocol.

### Preparation for the Semi-automated Processing

Installation of the macro software: For the semi-automated protocol, install Macro Scheduler software v15’ available at https://www.mjtnet.com.

**Note:** The user can use any image recognition and object-oriented macro software as preferred. Here, we describe the “Macro Scheduler 15” software as a reference.

### Step-by-step method details

#### Steps of Manual Processing

The following steps describe the conventional workflow for CM image analysis using the Imaris 10.1 software.

**Time: ∼60-90 minutes**

##### 1. Opening Image Files

a. Open the **Imaris** application.
b. In the toolbar, choose **‘Arena’** 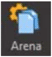**> ‘Observe Folder’** and choose the folder containing the image files converted in step 1 from the folder tree.
c. In the **‘Arena’** window, double click the image to be viewed.
d. In the **‘Surpass’** window 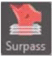, increase the **‘Rendering Quality’** to the highest setting (towards the right) from ‘**Volume**’ tab > under ‘**Scene**’ tree.
e. In the toolbar, choose **‘Image Processing’ > ‘Free Rotate**’ and change the **‘Angle’** setting to **<u>75º</u>** and click’OK’.
f. In the toolbar, choose **‘Edit’ > ‘Show Display Adjustment**’ to display the channel selections. The intensity of each channel can be changed by moving the histogram ranges found above and below each of the channel. Figure 1 shows an image of the computer screen after opening an image of a mouse cochlea in Imaris.

**Figure 1:**
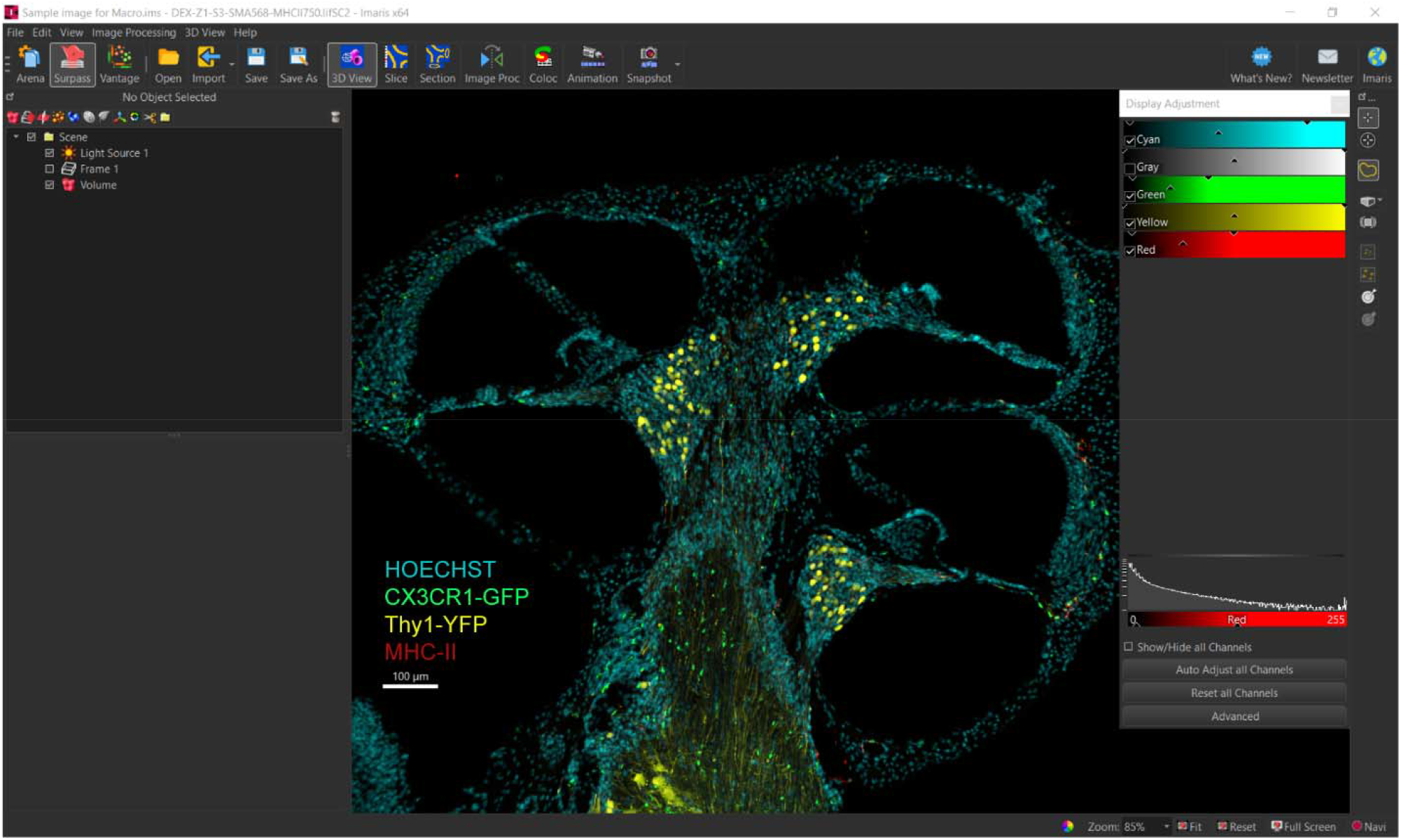
A representative image showing the computer screen after opening an image of a mouse cochlea in Imaris 10.1. The cross-sectional image present three cochlear turns with the modiolus placed centrally. Here, nuclei are stained with Hoechst 3342 (Thermofisher, Catalog# 62249, cyan). CX3CR1+ macrophages (green) and Thy1+ neurons (yellow) are labeled with enhanced green fluorescent protein (EGFP) and enhanced yellow fluorescent protein (eYFP) (yellow) respectively.

##### 2. Making Surfaces

a. In the ‘Scene’ tree, choose the **Surfaces tab** 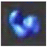 to make a surface for every area of interest.
b. Rename each surface to the corresponding identity of the region of interest by right-clicking the surface text.
c. Click the **‘Skip automatic creation, edit manually’** button. 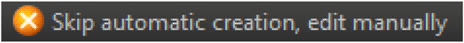
d. Choose the **Draw tab** 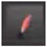 click the **‘Contour’** button then click the **‘Draw’** button. 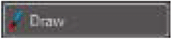
e. Using the brightfield channel, draw the outline of the region of interest by clicking the border. Remove all other channels by clicking the respective channels in the ‘Display Adjustment’ window.
f. Under Selection, click the **‘Copy’** button. Change the **‘Slice Position’** to the last slice and click the **‘Create Surface’** button.
g. Choose the **Edit tab** 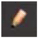. Under ‘Edit’, click the **‘Mask selection**..**’** button. In the Mask Channel Window, change the channel under ‘Channel Selection’, check ‘**Duplicate channel before applying mask’**, then click ‘OK’. Repeat for all channels, always masking the original channels.
h. Choose the **Statistics tab** 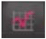, click the **‘Selection’** button, under the drop down ‘Specific Values’ list choose **‘Volume’** and record the value.

##### 3. Creating and Counting Spots

1. In the Display Adjustment’ window, unclick all channels except the masked channel to be analyzed.
2. Choose the Spots tab 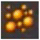, then click the blue forward arrow button 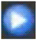 to change the page.
3. On page 2/4, under the drop down ‘Source Channel’ list choose the masked channel to be analyzed.
4. On page 2/4, change the ‘Estimated XY Diameter’ to **<u>7.58</u>** for macrophages and spiral ganglion neurons, or **<u>6.00</u>** for the nuclei. Click the blue forward arrow button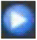.
5. On page 3/4, change the quality of the spot detection by moving the bottom histogram range left or right. Ensure the spots do not double count. Click the green double forward arrow button 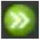 to finish.
6. Choose the **Edit tab** 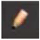 to manually count the additional cells. ‘Shift + click’ to count a cell. ‘Shift + click’ over a spot will un-count the cell.
7. Choose the **Statistics tab** 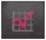 and click the **‘Overall’** button to see the number of spots counted.

However, while analyzing different specific regions of interest in one image, repetitive tasks such as the creation of surfaces, masks, and spots are needed.

#### Steps for developing the macro for semi-automated image analysis

**Time: ∼30 minutes**

##### 1. Creating the Surfaces

a. Creating configurations for surfaces for a single image:
  1. ‘Macro Scheduler 15’ was opened, and using the ‘New’ 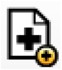 button, we created a Macro and saved it under the ‘SurfaceCreation’ (Supplementary S1). Figure 2 presents the opening window of Macro Scheduler 15.
  2. The ‘Object Recognition Wizard’ was used to crop the surface icon 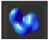, so the macro could click the surface icon as many surfaces as we intended to create (8 for the given example).
  3. Once the macro was consistently clicking the surface button, the ‘Tab’ button was implemented in code to toggle between each surface (example: going from Surface 1 to Surface 2), and keyboard commands were used to double-click the image with the F2 keystroke, and then enter the name of each surface.
  4. To subsequently set the resolution to maximum for each surface, the ‘Object Recognition Wizard’ was used to crop the ‘Manual’ box 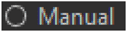, and the mouse was set to click the halfway point on the threshold bar 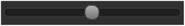, so that it could drag it to the right for maximum resolution. This was repeated for all 8 surfaces.
b. Creating configurations for surfaces for a batch of images:
  1. ‘Macro Scheduler 15’ was opened, and using the ‘New’ 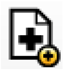 button, we created a Macro and saved it under the name ‘MasterSurface’ (Supplementary S2).
  2. The ‘Object Recognition Wizard’ was used to crop the ‘Close Tab’ option at the top right of the Imaris setup 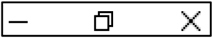, so the macro could click it after it applies the ‘SurfaceCreation’ to each file in the given folder.
  3. Next, we created variables ‘imarisExecutablePath’ and ‘copiedPath’, the current version of Imaris and the path of the desired folder, respectively. A for loop was created to iterate through each file in ‘copiedPath’ and run the ‘SurfaceCreation’ macro on each file.
  4. At the end of the macro, a wait time of 10 seconds was assigned to account for any differences between the computer’s frame rate and the macro’s operating rate. Then, the code for sending CTRL-S was implemented, and the code for clicking the ‘Close Tab’ option at the top right of the Imaris setup was created.

**Figure 2:**
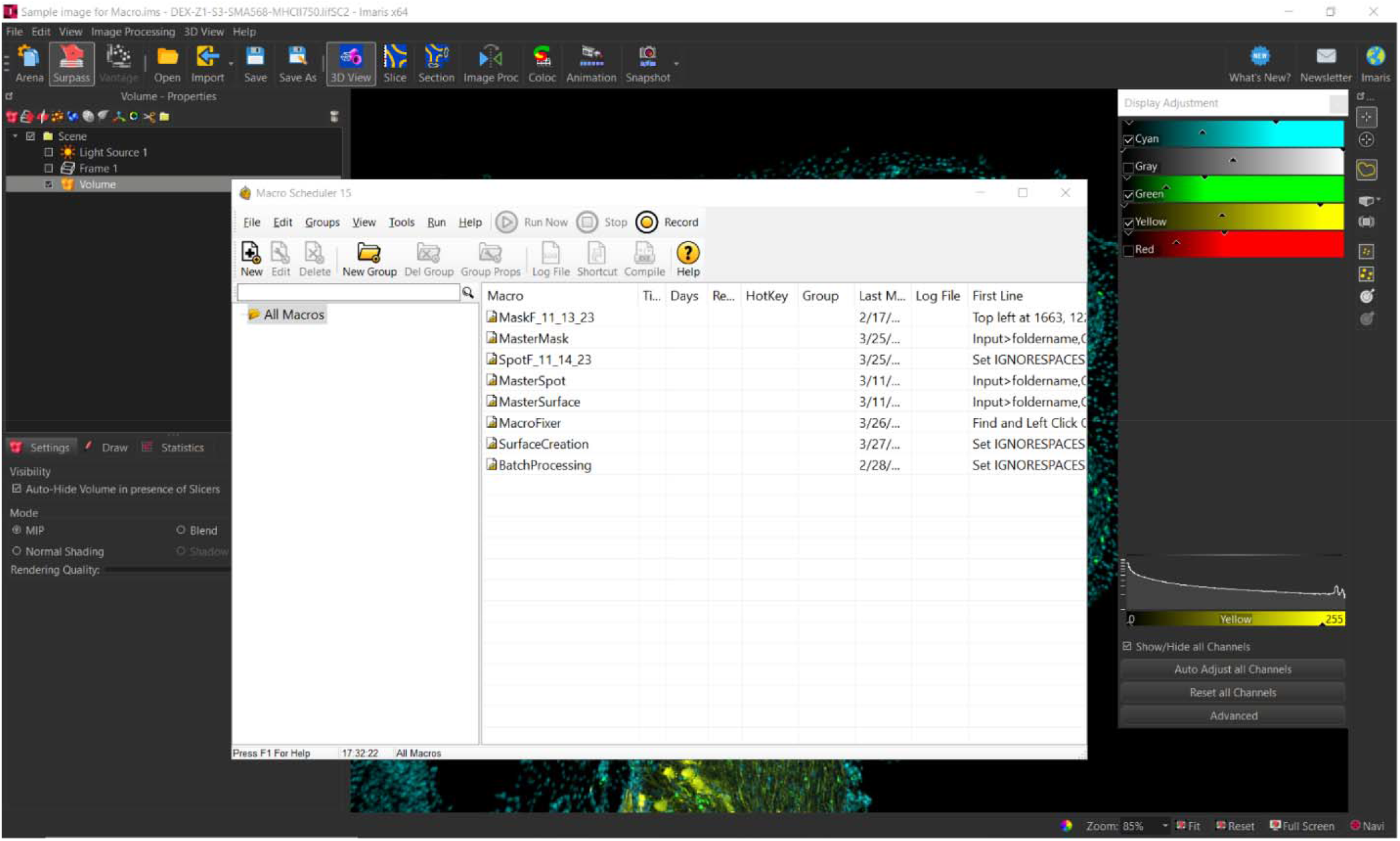
Opening window of Macro Scheduler 15.

##### 2. Creating masks for the surfaces

a. Creating Masks for the surfaces for an image:
  1. ‘Macro Scheduler 15’ was opened, and using the ‘New’ 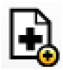 button, we created a Macro and saved it under the desired name (‘MaskF_11_13_23’ in our example, Supplementary S3).
  2. The ‘Object Recognition Wizard’ was used to crop the pencil for the ‘Edit’ button 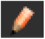, so the macro could click it 8 times (once per surface).
  3. The macro started by selecting the last surface, Modiolus, and setting it to the ‘Edit’ tab – after which keystrokes of the ‘Up Arrow’ were applied to traverse our surfaces in reverse order, setting all of them to the ‘Edit’ tab.
  4. Once the ‘ST’ was selected and set to the ‘Edit’ tab, we proceeded to implement code to create our first channel by using the ‘Object Recognition Wizard’ to crop the ‘Mask All’ button 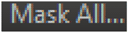 that was in the ‘Edit’ tab.
  5. After coding the macro to click on the ‘Mask All’, the ‘Object Recognition Wizard’ was used to crop the ‘Channel 1 – Cyan’ 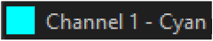 under the ‘Channel Selection’. A drop-down menu including all of our channels were available. The ‘Enter’ keystroke was coded so that the first channel, Nuclei ST, could be created.
  6. In the ‘Display Adjustment’ tab, there was a 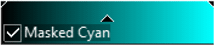 channel created after we selected the ‘Mask All’ button. Code was implemented to click on the ‘Masked Cyan’ text, which was followed by an ‘Image Properties’ tab.
  7. Subsequently, we coded up the functionality to click on the ‘Name’ space and send keystrokes for our desired channel, which is ‘Nuclei ST’. This process, from selecting ‘Mask All’ to naming the newly masked channel (whether it is blue, yellow, green, or red) was implemented for each surface. Note: Because of Imaris’s limitations on the amount of channels open, the macro was coded to click the ‘Show/Hide all channels’ 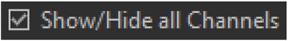 button in the ‘Display Adjustment tab, and this was unselected after the GM Nuclei mask channel was created.
b. Creating Masks for the surfaces for a batch of images:
  1. ‘Macro Scheduler 15’ was opened, and using the ‘New’ 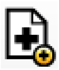 button, we created a Macro and saved it under the name ‘MasterMask’ for our purpose (Supplementary S4).
  2. The ‘Object Recognition Wizard’ was used to crop the ‘Close Tab’ option at the top right of the Imaris setup 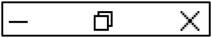, so the macro could click it after it applies the ‘MaskF_11_13_23’ to each file in the given folder.
  3. Next, variables ‘imarisExecutablePath’ and ‘copiedPath’, the current version of Imaris and the path of the desired folder respectively. A for loop was created to iterate through each file in ‘copiedPath’ and run the MaskF_11_13_23 macro on each file.
  4. At the end of the macro, a wait time of 10 seconds was assigned to account for any differences between the computer’s frame rate and the macro’s operating rate. Then, code for sending CTRL-S was implemented, and the code for clicking the ‘Close Tab’ option at the top right of the Imaris setup was created.

##### 3. Creating spots

a. Creating spot configurations for the surfaces for an image
  1. ‘Macro Scheduler 15’ was opened, and using the ‘New’ 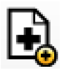 button, we created a Macro and saved it under the name ‘SpotF_11_14_23’ (Supplementary S5).
  2. The ‘Object Recognition Wizard’ was used to crop the yellow image of the spots in the left side of this image 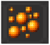, so the macro could click it 27 times (once per each mask channel).
  3. Once the macro was consistently clicking the spots button, the macro was coded to double click on the very last (27^th^) spot, and the F2 keystroke was implemented to name it in reverse order, Modiolus MHCII.
  4. We then traversed back up the spots, naming them with the F2 naming the spot and the ‘Up Arrow’ command toggling to the previous spot, until the macro arrived at the first spot, ‘ST Nuclei’. There, a ‘Create’ tab 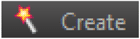 can be viewed, under which there is a ‘Skip automatic creation, edit manually’ option 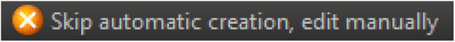, under which a blue arrow pointing right 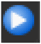 can be observed.
  5. Using the ‘Object Recognition Wizard’, we cropped the blue arrow pointing right 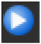 and the tab labelled ‘Source Channel’ is now able to be viewed.
  6. The macro selected the ‘Channel 1 – Cyan’ 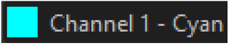 under the ‘Source Channel’ using the ‘Object Recognition Wizard’ to select the ST Nuclei mask channel.
  7. The macro traversed down to the ST Nuclei option in the drop-down menu using the ‘Down Arrow’ key for the necessary amounts of iterations. Then, the macro was programmed to press the ‘Enter’ keystroke. Under the ‘Spot Detection’ section, the macro clicked on the ‘Estimated XY Diameter’ and sent keystrokes with the correct values, depending on which type of mask channel (Nuclei is 6.00 and every other mask is 7.58 picometers).
  8. Once done, the Imaris screen should be on the threshold. The macro sequentially applied this to every spot, and the ‘ST Macrophage’ spot 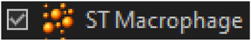 was selected using the ‘Object Recognition Wizard’ to traverse to other spots as a reference point (using ‘Down Arrow’ keystrokes to get to other spots).
b. Creating spot configurations for the surfaces for a batch of images:
  1. ‘Macro Scheduler 15’ was opened, and using the ‘New’ 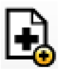 button, we created a Macro and saved it under the name ‘MasterSpot’ for our purpose (Supplementary S6).
  2. The ‘Object Recognition Wizard’ was used to crop the ‘Close Tab’ option at the top right of the Imaris setup 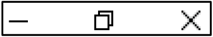, so the macro could click it after it applies the ‘SpotF_11_14_23to each file in the given folder.
  3. Next, variables ‘imarisExecutablePath’ and ‘copiedPath’, the current version of Imaris and the path of the desired folder respectively. A for loop was created to iterate through each file in ‘copiedPath’ and run the SpotF_11_14_23 macro on each file.
  4. At the end of the macro, a wait time of 10 seconds was assigned to account for any differences between the computer’s frame rate and the macro’s operating rate. Then, code for sending CTRL-S was implemented, and the code for clicking the ‘Close Tab’ option at the top right of the Imaris setup was created.

#### Steps for the semi-automated image analysis

We used our custom-made macros for semi-automated image analysis in following steps:

**Time: ∼30-45 minutes**

##### 1. Creating the Surfaces

a. Creating configurations for surfaces for a single image: After selecting the first macro, named ‘SurfaceCreation’, this macro will create all 8 surfaces, from scala tympani to Modiolus (for our protocol), and set the Resolution to ‘Manual’ and the maximum resolution possible for all 8 surfaces.
b. Creating configurations for surfaces for a batch of images: The Batch Processing macro, named ‘MasterSurface’, will apply the individual, object-recognition oriented, individual macro ‘SurfaceCreation’ to every single file in the desired folder. To run the ‘MasterSurface’ macro, there must not be any surfaces created, nor any spots or any masks (besides the default channels). The desired File Directory must be put inside of the ‘MasterSurface’ macro under the variable ‘CopiedPath’ (Figure 3).

**Figure 3:**
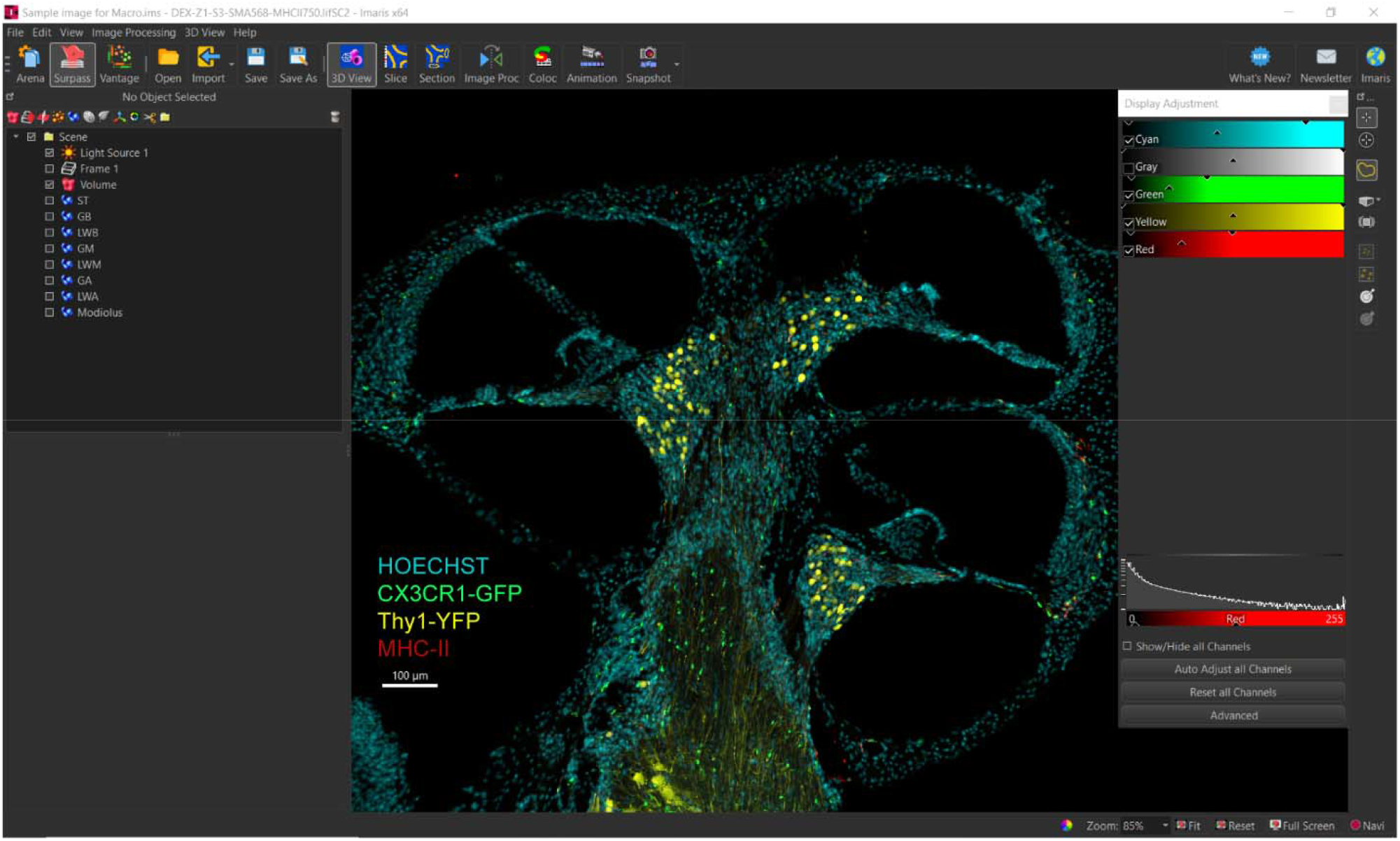
Output window after running the macro for naming surface. Using the Macro scheduler 15, a macro has been created that can open all the images in a folder sequentially and can perform the following functions a) Opening IMARIS image files, adjusting rendering quality, rotating, adjusting background, and intensity level of channels. b) Creating the names of the surfaces for all the images in the analysis folder.

**Note:** For the manual steps for creating surfaces, refer to steps 1-2 in the ‘Steps for the Manual Processing’ section of this paper.

##### 2. Drawing the Surface (Manual)

Refer to steps 3-6 in the ‘Protocol for the Manual Processing’ section of this paper, under the ‘Making Surfaces’ section (Figure 4).

**Figure 4:**
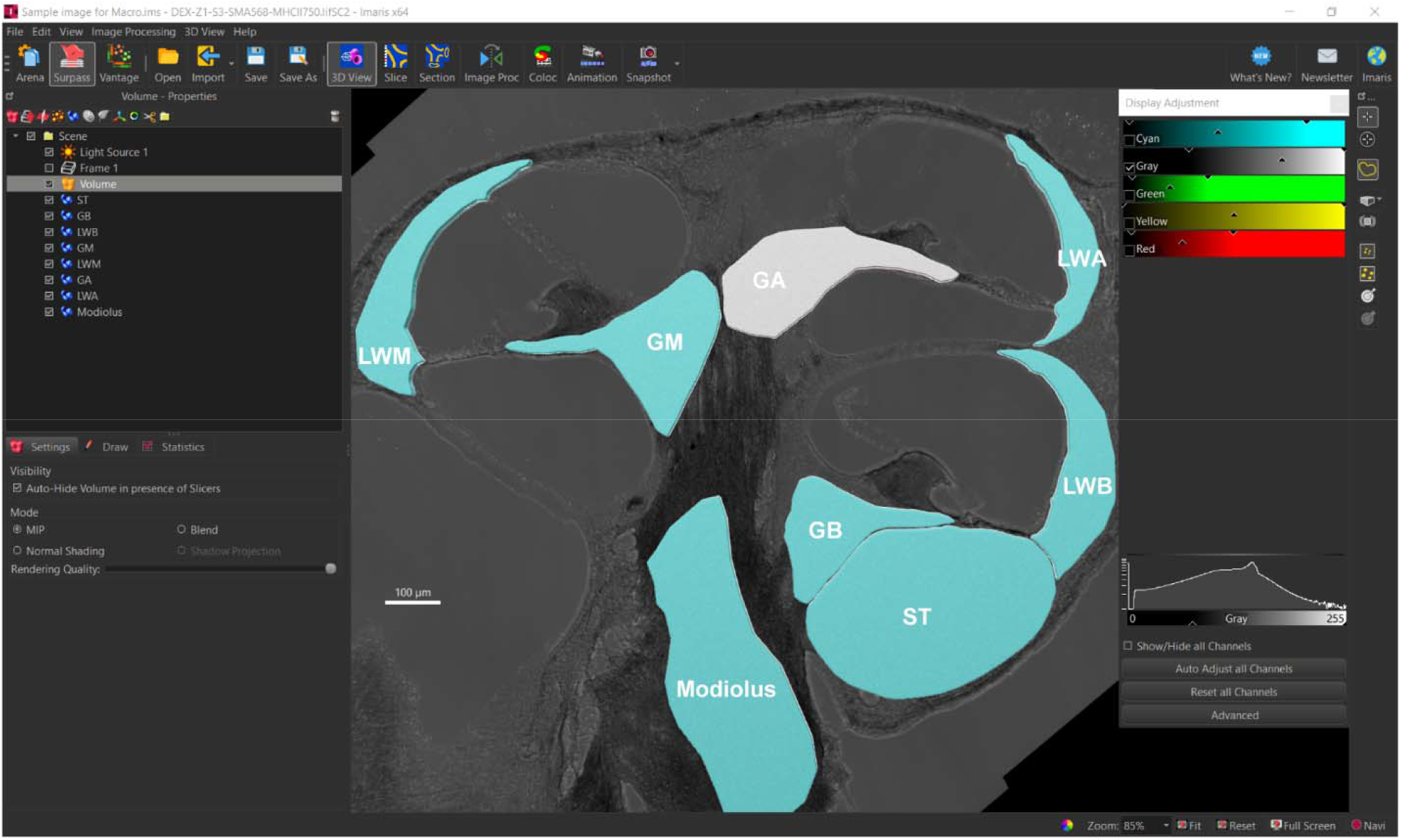
Manually drawn outlines of eight surfaces of cochlea. A total of 8 surfaces of the cochlea were drawn comprising of scala tympani of the base of the cochlea (ST), spiral ganglion of the base (GB), spiral ganglion of the middle (GM) and spiral ganglion of the apex (GA), lateral wall of the base (LWB), lateral wall of the middle (LWM), lateral wall of the apex (LWA) and Modiolus.

##### 3. Masking the Surface

a. Creating Masks for the surfaces for an image: After selecting the second macro, named ‘MaskF_11_13_23’, this macro will create all 33 masks for each surface, starting from scala tympani and ending with the Modiolus (for our protocol). The macro will go through the same procedure as the manual process, using object recognition to be precise; after going through the ‘Edit’ tab for each surface, it creates masks such as Nuclei, Macrophage, and MHCII (Neuron if applicable).
b. Creating Masks for the surfaces for a batch of images: The Batch Processing macro, named ‘MasterMask’, will apply the individual, object-recognition oriented, individual macro ‘MaskF_11_13_23’ to every single file in the desired folder. To run the ‘MasterMask’ macro, the surfaces must be created, the ‘Display Adjustment’ must have a checkmark next to ‘All Channels Selected’, and 4 channels must have individual check marks next to them. Additionally, the ‘Mask All’ option under the ‘Edit’ tab for each surface must be available (as shown by a light gray color when viewed). The user needs to put the File Directory of the user’s desired file inside of the ‘MasterMask’ macro under the variable ‘CopiedPath’ (Figure 5).

**Figure 5:**
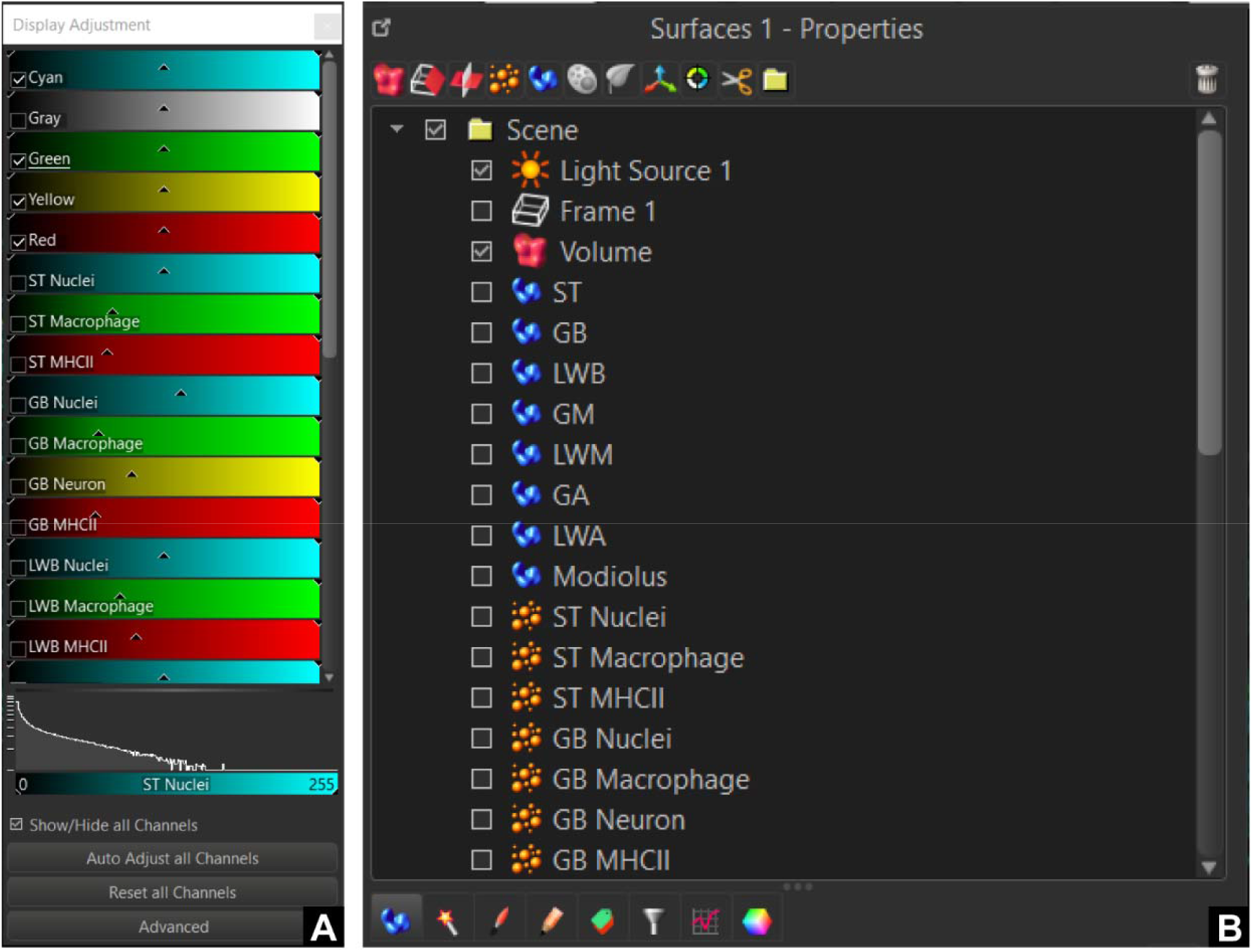
Automated creation of the masks for channels. Using a macro for mask creation, masks for every channel for each surface were created. The macro opens every image in the folder, creates masks for all of them, and saves the edited images. As we are analyzing 3 channels for 8 surfaces, a total number of 24 masks were created.

**Note:** For the manual steps for creating surfaces, refer to steps 1-2 in the ‘Steps for the Manual Processing’ section of this paper.

##### 4. Creating spots

a. Creating spot configurations for the surfaces for an image: After selecting the second macro, named ‘SpotF_11_13_23’, this macro will create all 33 spots for each surface, starting from scala tympani and ending with the Modiolus (for our protocol). The macro will go through the same procedure as the manual process, using object recognition to be precise; after going through the ‘Spot’ button 33 times, this macro names all the spots in chronological order, then goes individually to set the diameter as well as the corresponding mask for the spot.
b. Creating spot configurations for the surfaces for a batch of images: The Batch Processing macro, named ‘MasterSpot’, will apply the individual, object-recognition oriented, individual macro ‘SpotF_11_13_23’ to every single file in the desired folder. To run the ‘MasterSpot’ macro, there must not be any spots created, but the masks should be correctly created (and when a file is opened, it should give an Imaris error related to the graphic overload, as too many channels are selected), the ‘Display Adjustment’ tab should be open. The user need to put the File Directory of his/her desired file inside of the ‘MasterSpot’ macro under the variable ‘CopiedPath’.

**Note:** For the manual steps for creating spots, refer to steps in the ‘Creating and Counting Spots: of the ‘Steps for the Manual Processing’ section of this paper. The output spots for nucleus, neuron and macrophages are shown in Figure 6.

**Figure 6:**
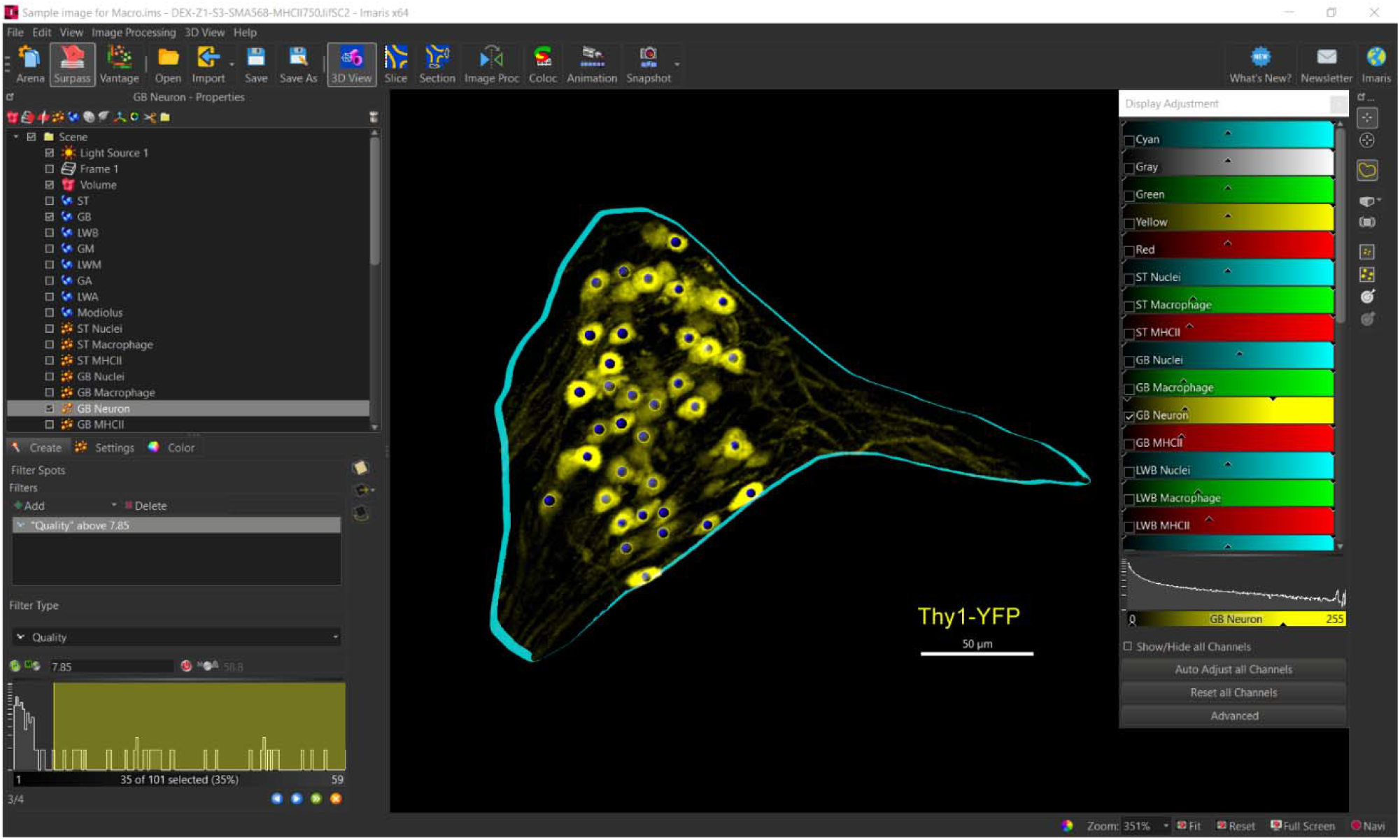
Automated creation of spots for each mask. Following selection of individual spot, the macro for spot selects predetermined parameters for creating the spots. The macro opens all images in the folder and creates spots for all the masks. The macro for spot creation does the task in following steps: a) Creation of the names of the spot b) Selection of the parameters for individual spots and create spots c) Creation of spots for all images in the analysis folder and saving using a macro. Thresholds for individual spots set at default value. The macro will sequentially create all the spots for individual images and for a batch as directed. The threshold is subject to manual modification.

##### 5. Setting the threshold and counting the Spots (Manual)

Under the ‘Create’ tab, the left margin of the slider is adjusted in accordance to ensure high specificity with minimal false positivity (Figure 7).

**Figure 7:**
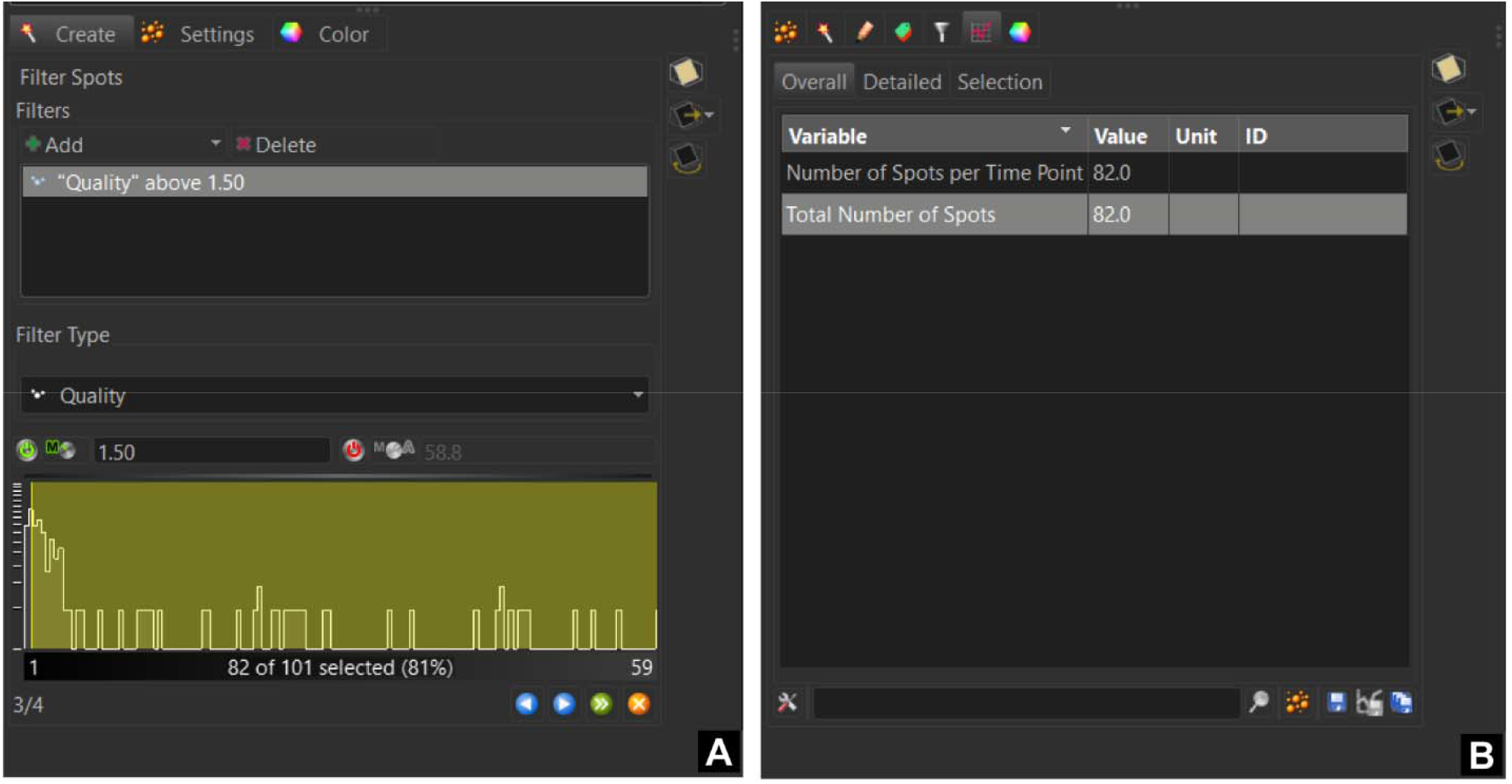
Manual thresholding for spots and logging data. The dynamic range for individual channels is manually adjusted using the Display Adjustment window. Threshold for the spots is manually adjusted to standards defined by the investigator(s). Additional ‘edit’ can be done using the edit option under the ‘spot’ tab. Window showing final output stage after running macros. **(A)** Here threshold for each spot is manually adjusted according to the pre-specified study requirement. **(B)** The data is collected or logged from the “Overall” or “Detailed” tab.

**Note:** Refer to steps in the ‘Creating and Counting Spots: of the ‘Steps for the Manual Processing’ section of this paper. The count then should be recorded.

**Critical:** As step 2 and 5 of this section of the paper is manual, hence we refer our macro to be ‘Semi-automated’.

Figure 8 shows an overview of the semi-automated workflow.

**Figure 8:**
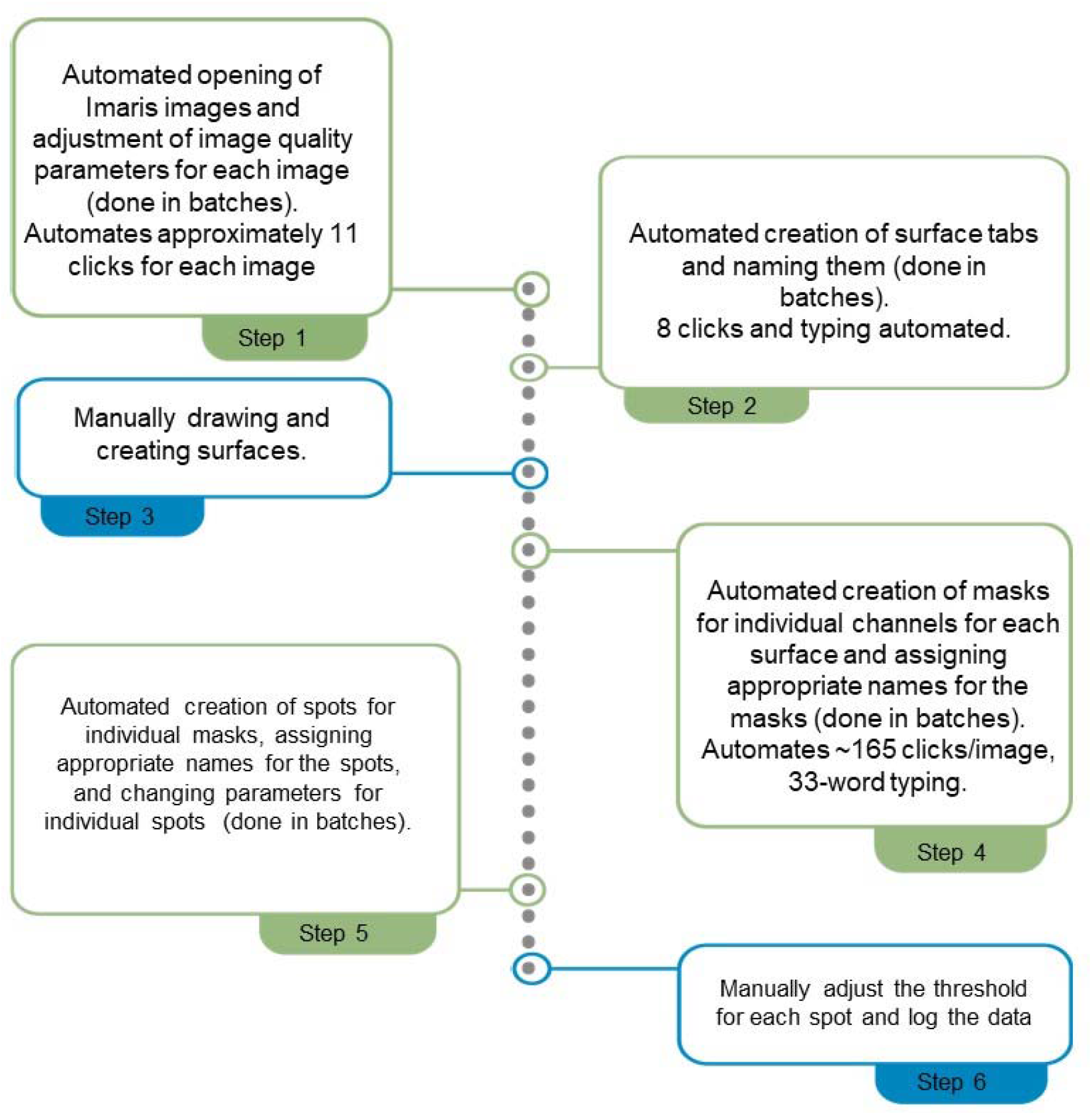
Overview of Semi-automated workflow for counting cells using Imaris and Macro-schedular software. By using Macro-scheduler software, 4 steps were automated in the counting process. Automated steps are shown in Green box and Blue indicates the steps that are were not automated. As steps 1,2,4 and 5 are done in batches the total time saved increases significantly.

### Expected outcomes

Our developed semi-automated macro tool can enhance efficiency by significantly reducing the manual labor performed by a researcher in image analysis tasks for biological research.

For example, manual creation and renaming a single surface with maximum resolution requires a six clicks per image. For creating eight surfaces in instance of the reference image in this article, a total of forty-eight clicks are required. Using the developed macro, this tedious process can be minimized to only one click through. By using the Macro-scheduler software, 4 out of 6 steps in the counting process has been automated.

Moreover, the macro introduces flexibility in terms of adapting to different devices. For instance, in the initially developed macro (Rahman, Khan, et al., 2023), a minimal change to the window size, predefined setup in any window or the arena workspace in the Imaris software would disrupt the workflow due to its dependence on a coordinate-based system of the device screen. As such, it lacked generalizability and intelligence and necessitated the development of a unique macro for a particular device. To address this, in the current version, ‘Image recognition’ and ‘object-oriented clicking’ features have been implemented resulting in the macro locating its object and clicking accordingly irrespective of the window and arena workspace. Consequently, the newly developed version of macro can be used on any compatible device by running the appropriate software.

Furthermore, the method not requiring any prior programming experience, can be reproduced by researchers working on cochlear samples or any other biological tissues from different disciplines. A study by Rahman and Mostaert, et al has implemented this semi-automated technique for mouse implanted cochlear samples (Rahman, Mostaert, et al., 2023). Additionally, the semi-automation should enhance consistency and reproducibility across samples, researchers within a lab, and across different labs. The integration of artificial intelligence and machine learning can potentially contribute to making it more efficient and user friendly.

### Limitations

Although the macro reduces time requirement for analysis of confocal microscope image dataset when run in batch, it does not significantly reduce the processing time when run for each image individually. As a solution, the latency time can be reduced by increasing the running speed in the recorded macro code. However, one drawback is that, running the macro at a speed higher than 1x can occasionally make it unstable due to the computer system’s limitation in processing a large amount of data at a time. This latency requires to be taken into account when setting the run speed for the macro.

### Troubleshooting

#### Problem 1

Differences between the computer and macro’s frame rate (corresponds to step 6b of “Steps for developing the Semi-automated Processing”).

The macro can speed faster than the computer processing due to computer lag arising from a multitude of reasons, including the workload on the computer chip, prolonged period of uninterrupted running, etc. This mismatch in speed can result in the macro clicking in unwanted places leading to erroneous results. Below are solutions for your considerations, should the macro be operating on a computer having speed lower than macro processing speed.

#### Potential solution

- Restart the computer: Restarting the computer and waiting ∼10 minutes can significantly reduce the amount of lag present in the computer. This is because the computer ceases all existing operations once it restarts.
- Shut down the computer: Shutting down the computer and starting it again when the user needs to use the macro (with a sufficient gap in time) can significantly reduce the amount of lag present in the computer. This is a recommended practice and makes it very likely that the computer and the macro have the same frame rate.

#### Problem 2

The macro starts to glitch, manifested by dark patches in the “Macro Scheduler 15” tab, and subsequently present in any macro you open. Also characterized by messages in the “Macro Scheduler 15” tab that mention “Insufficient memory resources”.

#### Potential solution

- Restart the computer: Restarting the computer and waiting ∼10 minutes can significantly reduce the amount of lag present in the computer. This is because the computer ceases all existing operations once it restarts.
- Shut down the computer: Shutting down the computer and starting it again when the user needs to use the macro (with a sufficient gap in time) can significantly reduce the amount of lag present in the computer. This is a recommended practice and makes it very likely that the computer and the macro have the same frame rate.

## Supporting information

Supplementary S3: Codes for Macro necessary for creation of masks

Supplementary S4: Codes for Macro necessary for creation of masks in a batch

Supplementary S6: Codes for Macro necessary for creation of spots in a batch

Supplementary S2: Codes for Macro necessary for creation of surface in a batch

Supplementary S5: Codes for Macro necessary for creation of spots

Supplementary S1: Codes for Macro necessary for creation of surface

## Acknowledgments

The research was supported by National Institute of Deafness and other Communication Disorders (NIDCD) R01 DC018488 (MH), R01 DC012578 (MH), P50 DC000242 (MH), F32 DC020643 (MR), National Center for Advancing Translational Sciences (NCATS) UM1TR004403 (MH), Cochlear corporation (MH), Alpha Omega Alpha Carolyn L. Kuckein Student Research Fellowship (PE), Iowa MSTP SUMR program (BH) and Pomerantz Career Center Hawkeye Research Experience Grant (BH).

## Author contributions

Conceptualization: MTR, MRH and NAK; Experiment: MTR, NAK, CG, IR, PE, SMF; Data analysis: MTR, CG, IR, PE, SMF; Figures: IR, MTR, SSL, MRH; Manuscript writing: MTR, MIM, NAK, SSL, IR, MRH; Manuscript review: All authors; Funding acquisition: MRH, MTR.

**Supplementary S1: Codes for Macro necessary for creation of surface: ‘SurfaceCreation’**.

**Supplementary S2: Codes for Macro necessary for creation of surface in a batch: ‘MasterSurface’**.

**Supplementary S3: Codes for Macro necessary for creation of masks: ‘MaskF_11_13_23’**.

**Supplementary S4: Codes for Macro necessary for creation of masks in a batch: ‘Master_Mask’**.

**Supplementary S5: Codes for Macro necessary for creation of spots: ‘SpotF_11_14_23’**.

**Supplementary S6: Codes for Macro necessary for creation of spots in a batch: ‘Master_Spot’**.

