## Supplementary S1: Codes for Macro necessary for creation of surface for "Supervised Automation of Cell Counting in Confocal Microscopic Cochlear Imaging Datasets Using Macro in Imaris"

//Set IGNORESPACES to 1 to force script interpreter to ignore spaces.
//If using IGNORESPACES quote strings in {" ... "}
//Let>IGNORESPACES=1


Let>spotCnt=7+1
Let>k=0
  Repeat>k
  //Find and Left Click Center of 
  FindImagePos>%BMP_DIR%\image_1.bmp,SCREEN,0.7,8,XArr,YArr,NumFound,CCOEFF
  If>NumFound>0
    MouseMove>XArr_0,YArr_0
    LClick
  Endif
  Let>k=k+1
Until>k,spotCnt

Wait>2
//Find and Left Click Center of 
FindImagePos>%BMP_DIR%\image_22.bmp,SCREEN,0.7,1,XArr,YArr,NumFound,CCOEFF
If>NumFound>0
  Wait>1
  MouseMove>XArr_0,YArr_0
  LDblClick
Endif


Wait>1

Send>Modiolus
Press Enter
Press Up
Press F2
Send>LWA
Press Enter
Press Up
Press F2
Send>GA
Press Enter
Press Up
Press F2
Send>LWM
Press Enter
Press Up
Press F2
Send>GM
Press Enter
Press Up
Press F2
Send>LWB
Press Enter
Press Up
Press F2
Send>GB
Press Enter
Press Up
Press F2
Send>ST
Press Enter

Wait>2

//Find and Left Click Center of 
FindImagePos>%BMP_DIR%\image_20.bmp,SCREEN,0.7,1,XArr,YArr,NumFound,CCOEFF
If>NumFound>0
  Wait>0.5
  MouseMove>XArr_0,YArr_0
  LClick
Endif

Wait>2
//Find and Left Click Middle Bottom of 
FindImagePos>%BMP_DIR%\image_16.bmp,SCREEN,0.7,6,XArr,YArr,NumFound,CCOEFF
If>NumFound>0
  Wait>0.5
  MouseMove>XArr_0,YArr_0
  LClick
Endif

Wait>2

//Find and Left Click Center of 
FindImagePos>%BMP_DIR%\image_18.bmp,SCREEN,0.7,1,XArr,YArr,NumFound,CCOEFF
If>NumFound>0
  Wait>0.5
  MouseMove>XArr_0,YArr_0
Endif

Wait>2

//Find and Left Click Center of 
FindImagePos>%BMP_DIR%\image_5.bmp,SCREEN,0.7,1,XArr,YArr,NumFound,CCOEFF
If>NumFound>0
  Wait>0.5
  MouseMove>XArr_0,YArr_0
  LDown
Endif

Wait>2

//Find and Left Click Center of 
FindImagePos>%BMP_DIR%\image_6.bmp,SCREEN,0.7,1,XArr,YArr,NumFound,CCOEFF
If>NumFound>0
  Wait>0.5
  MouseMove>XArr_0,YArr_0
  LUp
Endif

Wait>2

//Find and Left Click Center of 
FindImagePos>%BMP_DIR%\image_7.bmp,SCREEN,0.7,1,XArr,YArr,NumFound,CCOEFF
If>NumFound>0
  MouseMove>XArr_0,YArr_0
  LClick
Endif

Wait>2

//Find and Left Click Center of 
FindImagePos>%BMP_DIR%\image_20.bmp,SCREEN,0.7,1,XArr,YArr,NumFound,CCOEFF
If>NumFound>0
  Wait>0.5
  MouseMove>XArr_0,YArr_0
  LClick
Endif

Wait>2
//Find and Left Click Middle Bottom of 
FindImagePos>%BMP_DIR%\image_16.bmp,SCREEN,0.7,6,XArr,YArr,NumFound,CCOEFF
If>NumFound>0
  Wait>0.5
  MouseMove>XArr_0,YArr_0
  LClick
Endif

Wait>2

//Find and Left Click Center of 
FindImagePos>%BMP_DIR%\image_18.bmp,SCREEN,0.7,1,XArr,YArr,NumFound,CCOEFF
If>NumFound>0
  Wait>0.5
  MouseMove>XArr_0,YArr_0
Endif

Wait>2

//Find and Left Click Center of 
FindImagePos>%BMP_DIR%\image_5.bmp,SCREEN,0.7,1,XArr,YArr,NumFound,CCOEFF
If>NumFound>0
  Wait>0.5
  MouseMove>XArr_0,YArr_0
  LDown
Endif

Wait>2

//Find and Left Click Center of 
FindImagePos>%BMP_DIR%\image_6.bmp,SCREEN,0.7,1,XArr,YArr,NumFound,CCOEFF
If>NumFound>0
  Wait>0.5
  MouseMove>XArr_0,YArr_0
  LUp
Endif

Wait>2

//Find and Left Click Center of 
FindImagePos>%BMP_DIR%\image_8.bmp,SCREEN,0.7,1,XArr,YArr,NumFound,CCOEFF
If>NumFound>0
  MouseMove>XArr_0,YArr_0
  LClick
Endif

Wait>2

//Find and Left Click Center of 
FindImagePos>%BMP_DIR%\image_20.bmp,SCREEN,0.7,1,XArr,YArr,NumFound,CCOEFF
If>NumFound>0
  Wait>0.5
  MouseMove>XArr_0,YArr_0
  LClick
Endif

Wait>2
//Find and Left Click Middle Bottom of 
FindImagePos>%BMP_DIR%\image_16.bmp,SCREEN,0.7,6,XArr,YArr,NumFound,CCOEFF
If>NumFound>0
  Wait>0.5
  MouseMove>XArr_0,YArr_0
  LClick
Endif

Wait>2

//Find and Left Click Center of 
FindImagePos>%BMP_DIR%\image_18.bmp,SCREEN,0.7,1,XArr,YArr,NumFound,CCOEFF
If>NumFound>0
  Wait>0.5
  MouseMove>XArr_0,YArr_0
Endif

Wait>2

//Find and Left Click Center of 
FindImagePos>%BMP_DIR%\image_5.bmp,SCREEN,0.7,1,XArr,YArr,NumFound,CCOEFF
If>NumFound>0
  Wait>0.5
  MouseMove>XArr_0,YArr_0
  LDown
Endif

Wait>2

//Find and Left Click Center of 
FindImagePos>%BMP_DIR%\image_6.bmp,SCREEN,0.7,1,XArr,YArr,NumFound,CCOEFF
If>NumFound>0
  Wait>0.5
  MouseMove>XArr_0,YArr_0
  LUp
Endif

Wait>2

//Find and Left Click Center of 
FindImagePos>%BMP_DIR%\image_9.bmp,SCREEN,0.7,1,XArr,YArr,NumFound,CCOEFF
If>NumFound>0
  MouseMove>XArr_0,YArr_0
  LClick
Endif

Wait>2

//Find and Left Click Center of 
FindImagePos>%BMP_DIR%\image_20.bmp,SCREEN,0.7,1,XArr,YArr,NumFound,CCOEFF
If>NumFound>0
  Wait>0.5
  MouseMove>XArr_0,YArr_0
  LClick
Endif

Wait>2
//Find and Left Click Middle Bottom of 
FindImagePos>%BMP_DIR%\image_16.bmp,SCREEN,0.7,6,XArr,YArr,NumFound,CCOEFF
If>NumFound>0
  Wait>0.5
  MouseMove>XArr_0,YArr_0
  LClick
Endif

Wait>2

//Find and Left Click Center of 
FindImagePos>%BMP_DIR%\image_18.bmp,SCREEN,0.7,1,XArr,YArr,NumFound,CCOEFF
If>NumFound>0
  Wait>0.5
  MouseMove>XArr_0,YArr_0
Endif

Wait>2

//Find and Left Click Center of 
FindImagePos>%BMP_DIR%\image_5.bmp,SCREEN,0.7,1,XArr,YArr,NumFound,CCOEFF
If>NumFound>0
  Wait>0.5
  MouseMove>XArr_0,YArr_0
  LDown
Endif

Wait>2

//Find and Left Click Center of 
FindImagePos>%BMP_DIR%\image_6.bmp,SCREEN,0.7,1,XArr,YArr,NumFound,CCOEFF
If>NumFound>0
  Wait>0.5
  MouseMove>XArr_0,YArr_0
  LUp
Endif

Wait>2

//Find and Left Click Center of 
FindImagePos>%BMP_DIR%\image_10.bmp,SCREEN,0.7,1,XArr,YArr,NumFound,CCOEFF
If>NumFound>0
  MouseMove>XArr_0,YArr_0
  LClick
Endif

Wait>2

//Find and Left Click Center of 
FindImagePos>%BMP_DIR%\image_20.bmp,SCREEN,0.7,1,XArr,YArr,NumFound,CCOEFF
If>NumFound>0
  Wait>0.5
  MouseMove>XArr_0,YArr_0
  LClick
Endif

Wait>2
//Find and Left Click Middle Bottom of 
FindImagePos>%BMP_DIR%\image_16.bmp,SCREEN,0.7,6,XArr,YArr,NumFound,CCOEFF
If>NumFound>0
  Wait>0.5
  MouseMove>XArr_0,YArr_0
  LClick
Endif

Wait>2

//Find and Left Click Center of 
FindImagePos>%BMP_DIR%\image_18.bmp,SCREEN,0.7,1,XArr,YArr,NumFound,CCOEFF
If>NumFound>0
  Wait>0.5
  MouseMove>XArr_0,YArr_0
Endif

Wait>2

//Find and Left Click Center of 
FindImagePos>%BMP_DIR%\image_5.bmp,SCREEN,0.7,1,XArr,YArr,NumFound,CCOEFF
If>NumFound>0
  Wait>0.5
  MouseMove>XArr_0,YArr_0
  LDown
Endif

Wait>2

//Find and Left Click Center of 
FindImagePos>%BMP_DIR%\image_6.bmp,SCREEN,0.7,1,XArr,YArr,NumFound,CCOEFF
If>NumFound>0
  Wait>0.5
  MouseMove>XArr_0,YArr_0
  LUp
Endif

Wait>2

//Find and Left Click Center of 
FindImagePos>%BMP_DIR%\image_11.bmp,SCREEN,0.7,1,XArr,YArr,NumFound,CCOEFF
If>NumFound>0
  MouseMove>XArr_0,YArr_0
  LClick
Endif

Wait>2

//Find and Left Click Center of 
FindImagePos>%BMP_DIR%\image_20.bmp,SCREEN,0.7,1,XArr,YArr,NumFound,CCOEFF
If>NumFound>0
  Wait>0.5
  MouseMove>XArr_0,YArr_0
  LClick
Endif

Wait>2
//Find and Left Click Middle Bottom of 
FindImagePos>%BMP_DIR%\image_16.bmp,SCREEN,0.7,6,XArr,YArr,NumFound,CCOEFF
If>NumFound>0
  Wait>0.5
  MouseMove>XArr_0,YArr_0
  LClick
Endif

Wait>2

//Find and Left Click Center of 
FindImagePos>%BMP_DIR%\image_18.bmp,SCREEN,0.7,1,XArr,YArr,NumFound,CCOEFF
If>NumFound>0
  Wait>0.5
  MouseMove>XArr_0,YArr_0
Endif

Wait>2

//Find and Left Click Center of 
FindImagePos>%BMP_DIR%\image_5.bmp,SCREEN,0.7,1,XArr,YArr,NumFound,CCOEFF
If>NumFound>0
  Wait>0.5
  MouseMove>XArr_0,YArr_0
  LDown
Endif

Wait>2

//Find and Left Click Center of 
FindImagePos>%BMP_DIR%\image_6.bmp,SCREEN,0.7,1,XArr,YArr,NumFound,CCOEFF
If>NumFound>0
  Wait>0.5
  MouseMove>XArr_0,YArr_0
  LUp
Endif

Wait>2

//Find and Left Click Center of 
FindImagePos>%BMP_DIR%\image_12.bmp,SCREEN,0.7,1,XArr,YArr,NumFound,CCOEFF
If>NumFound>0
  MouseMove>XArr_0,YArr_0
  LClick
Endif

Wait>2

//Find and Left Click Center of 
FindImagePos>%BMP_DIR%\image_20.bmp,SCREEN,0.7,1,XArr,YArr,NumFound,CCOEFF
If>NumFound>0
  Wait>0.5
  MouseMove>XArr_0,YArr_0
  LClick
Endif

Wait>2
//Find and Left Click Middle Bottom of 
FindImagePos>%BMP_DIR%\image_16.bmp,SCREEN,0.7,6,XArr,YArr,NumFound,CCOEFF
If>NumFound>0
  Wait>0.5
  MouseMove>XArr_0,YArr_0
  LClick
Endif

Wait>2

//Find and Left Click Center of 
FindImagePos>%BMP_DIR%\image_18.bmp,SCREEN,0.7,1,XArr,YArr,NumFound,CCOEFF
If>NumFound>0
  Wait>0.5
  MouseMove>XArr_0,YArr_0
Endif

Wait>2

//Find and Left Click Center of 
FindImagePos>%BMP_DIR%\image_5.bmp,SCREEN,0.7,1,XArr,YArr,NumFound,CCOEFF
If>NumFound>0
  Wait>0.5
  MouseMove>XArr_0,YArr_0
  LDown
Endif

Wait>2

//Find and Left Click Center of 
FindImagePos>%BMP_DIR%\image_6.bmp,SCREEN,0.7,1,XArr,YArr,NumFound,CCOEFF
If>NumFound>0
  Wait>0.5
  MouseMove>XArr_0,YArr_0
  LUp
Endif

Wait>2

//Find and Left Click Center of 
FindImagePos>%BMP_DIR%\image_13.bmp,SCREEN,0.7,1,XArr,YArr,NumFound,CCOEFF
If>NumFound>0
  MouseMove>XArr_0,YArr_0
  LClick
Endif

Wait>2

//Find and Left Click Center of 
FindImagePos>%BMP_DIR%\image_20.bmp,SCREEN,0.7,1,XArr,YArr,NumFound,CCOEFF
If>NumFound>0
  Wait>0.5
  MouseMove>XArr_0,YArr_0
  LClick
Endif

Wait>2
//Find and Left Click Middle Bottom of 
FindImagePos>%BMP_DIR%\image_16.bmp,SCREEN,0.7,6,XArr,YArr,NumFound,CCOEFF
If>NumFound>0
  Wait>0.5
  MouseMove>XArr_0,YArr_0
  LClick
Endif

Wait>2

//Find and Left Click Center of 
FindImagePos>%BMP_DIR%\image_18.bmp,SCREEN,0.7,1,XArr,YArr,NumFound,CCOEFF
If>NumFound>0
  Wait>0.5
  MouseMove>XArr_0,YArr_0
Endif

Wait>2

//Find and Left Click Center of 
FindImagePos>%BMP_DIR%\image_5.bmp,SCREEN,0.7,1,XArr,YArr,NumFound,CCOEFF
If>NumFound>0
  Wait>0.5
  MouseMove>XArr_0,YArr_0
  LDown
Endif

Wait>2

//Find and Left Click Center of 
FindImagePos>%BMP_DIR%\image_6.bmp,SCREEN,0.7,1,XArr,YArr,NumFound,CCOEFF
If>NumFound>0
  Wait>0.5
  MouseMove>XArr_0,YArr_0
  LUp
Endif

Wait>2
